# Swarmer cell development of the bacterium *Proteus mirabilis* requires the conserved ECA biosynthesis gene, *rffG*

**DOI:** 10.1101/198622

**Authors:** Kristin Little, Murray J. Tipping, Karine A. Gibbs

## Abstract

Individual cells of the bacterium *Proteus mirabilis* can elongate up to 40-fold on surfaces before engaging in a cooperative surface-based motility termed swarming. How cells regulate this dramatic morphological remodeling remains an open question. In this paper, we move forward the understanding of this regulation by demonstrating that *P. mirabilis* requires the gene *rffG* for swarmer cell elongation and subsequent swarm motility. The *rffG* gene encodes a protein homologous to the dTDP-glucose 4,6 dehydratase protein of *Escherichia coli*, which contributes to Enterobacterial Common Antigen biosynthesis. Here we characterize the *rffG* gene in *P. mirabilis*, demonstrating that it is required for the production of large lipopolysaccharide-linked moieties necessary for wild-type cell envelope integrity. We show that absence of the *rffG* gene induces several stress-responsive pathways including those controlled by the transcriptional regulators RpoS, CaiF, and RcsB. We further show that in *rffG*-deficient cells, suppression of the Rcs phosphorelay, via loss of RcsB, is sufficient to induce cell elongation and swarm motility. However, loss of RcsB does not rescue cell envelope integrity defects and instead results in abnormally shaped cells, including cells producing more than two poles. We conclude that a RcsB-mediated response acts to suppress emergence of shape defects in cell envelope-compromised cells, suggesting an additional role for RcsB in maintaining cell morphology under stress conditions. We further propose that the composition of the cell envelope acts as a checkpoint before cells initiate swarmer cell elongation and motility.

**Importance statement:** *P. mirabilis* swarm motility has been implicated in pathogenesis. We have found that cells deploy multiple uncharacterized strategies to handle cell envelope stress beyond the Rcs phosphorelay when attempting to engage in swarm motility. While RcsB is known to directly inhibit the master transcriptional regulator for swarming, we have shown an additional role for RcsB in protecting cell morphology. These data support a growing appreciation that the Rcs phosphorelay is a multi-functional regulator of cell morphology in addition to its role in microbial stress responses. These data also strengthen the paradigm that outer membrane composition is a crucial checkpoint for modulating entry into swarm motility. Furthermore, the *rffG*-dependent moieties provide a novel, attractive target for potential antimicrobials.

## Introduction

Bacteria can migrate across a surface using a cooperative group motility termed swarming. For *Proteus mirabilis*, a gram-negative opportunistic pathogen, rapid surface-based swarm motility likely contributes to its pathogenesis during catheter associated urinary tract infections (1, 2). On hard agar (1.5 - 2%) surfaces, cells elongate from ~ 2 µm rods into hyper-flagellated, snake-like "swarmer" cells that carry multiple chromosomes and range in length from 10 – 80 µm (3-5). Cell elongation and enhanced flagellar gene expression are considered genetically linked and occur upon growth on a hard agar surface (6-9). Multiple swarmer cells closely associate into rafts that collectively move across a surface (3, 5, 10). After a defined period of motility, swarmer cells divide into short (1 – 2 µm) non-motile rod-shaped cells (11). Iterative rounds of swarmer cell elongation, group motility, and cell division comprise the swarmer cell developmental cycle and result in the rapid occupation of centimeter-scale surfaces in a stereotypical concentric ring pattern (11).

Swarmer cell elongation entails a broad range of physiological changes in addition to the dramatic morphological remodeling. Dozens of genes experience drastic changes in expression; for example, genes for flagella production become up-regulated on surfaces (12-16). Elongation into swarmer cells is also coordinated with changes to the cell envelope that minimally include alterations of the outer membrane. For example, in the outer membrane, lipopolysaccharide (LPS) structure is modified, fluidity is increased, and areas of phospholipid bilayer arise (17-19). Cells also transition through unknown mechanisms from being rigid to flexible (3-5).

In many bacteria, a bidirectional relationship exists between swarmer cell motility and cell envelope structure. There are several cell envelope biosynthesis pathways in Enterobacteriaceae such as LPS and Enterobacterial Common Antigen (ECA) biosynthesis for the outer membrane and peptidoglycan biosynthesis for the cell wall. Interrogating the specific contribution of each pathway to *P. mirabilis* swarmer cell development has proven challenging, partly because these three pathways share a pool of substrates (20, 21). For example, in *Escherichia coli*, genetic modifications to each of these biosynthetic pathways can dramatically alter cell shape and motility due to perturbations in the balance of the shared cell envelope substrate, undecaprenyl phosphate (20, 21). Moreover, disruption of cell envelope-associated genes inhibits swarmer cell development and motility of *P. mirabilis* through several mechanisms. For example, loss of the LPS biosynthesis gene *waaL* (22, 23) inhibits swarmer cell elongation and motility through activation of the Rcs phosphorelay (23), while the stress-associated sigma factor RpoE (24, 25) responds to disruptions of the LPS biosynthesis gene *ugd*(25). Less is known about role of ECA biosynthesis in *P. mirabilis*.

Cell envelope structure and stress sensing also appear to play broadly conserved roles in the swarm regulation of many bacterial species, including *P. mirabilis*, *E. coli*, and *Serratia marcescens* (20-22, 24, 26-30). In the aforementioned organisms, the Rcs (regulator of capsule synthesis) phosphorelay, which is a complex cell envelope stress-sensing signal transduction pathway, plays a key role in swarm motility inhibition (22, 26, 31). The Rcs phosphorelay, through the transcriptional regulator RcsB, directly represses the *flhDC* genes, which themselves encode the master transcriptional regulator of swarming, FlhD_4_C_2_ (27, 29). The current paradigm is that cell envelope stress or outer membrane defects activate membrane-localized Rcs proteins, which then phosphorylate and activate the response regulator RcsB (22, 26, 27, 31) (see also reviews (32, 33)). Decreased levels of *flhDC* result in reduced flagella production and the failure of cells to elongate, thus inhibiting swarm motility. RcsB directly activates the expression of the cell division-related genes, *minCDE*; however, the molecular mechanisms of this regulation remain unclear (6, 7). RcsB also induces production of several fimbrial genes, including paralogues of the fimbrial transcriptional regulator MrpJ. Together, RcsB and MrpJ modulate broad transcriptional and behavioral changes to promote cell adherence and biofilm formation and to repress swarm motility (7, 34).

Here we address the role of the cell envelope and stress sensing pathways in the regulation of swarmer cell development, an early stage of swarm motility. We show that *P. mirabilis* cells require the *rffG* gene, which is predicted to encode the sugar-modifying enzyme dTDP glucose-4,6-dehydratase, to produce an uncharacterized LPS-linked structural component of the cell envelope. As a homologous *rffG* gene and its conserved cluster of flanking genes are responsible for ECA production in *E. coli* (35), we posit that these structures may be ECA-derived. We further show that cells lacking the *rffG* gene remain short on swarm-permissive surfaces and suffer from cell envelope integrity defects that make elongated cells more susceptible to rupturing. We found that *rffG-*dependent moieties were not physically required for swarmer cell elongation; instead, loss of the *rffG* gene activated several swarm-inhibitory pathways, including the Rcs phosphorelay. Indeed, a RcsB-mediated response was sufficient to restrict swarmer cell elongation of *rffG*-deficient cells by inhibiting *flhDC* expression. We have also identified a novel role for RcsB in the maintenance of cell morphology during swarmer cell elongation. We found that RcsB was necessary to suppress the cell morphology of *rffG*-deficient cells that were genetically forced to elongate into swarmer cells. We posit that cell envelope composition is a crucial signaling checkpoint before entry into surface-based swarm motility. The Rcs phosphorelay response regulator not only mediates this signaling checkpoint, but also serves an important role in maintaining a normal cell shape during swarmer cell elongation.

## Results

### Cells require the *rffG* gene to complete swarmer cell elongation and initiate swarming

Previous research has explored the role of LPS biosynthesis genes in the regulation of *P. mirabilis* swarm motility, but a role for ECA has not been described (23, 25). Here, we interrogated the role in swarming of a gene associated with ECA biosynthesis. We characterized a swarm-deficient mutant strain presumably incapable of producing ECA by generating a chromosomal deletion of the *rffG* gene in *P. mirabilis* strain BB2000, resulting in a Δ*rffG* strain. A colony of the wild-type strain occupied a circle of 10-centimeter diameter by 24 hours on swarm-permissive and nutrient-rich CM55 agar; however, colonies of the Δ*rffG* population did not expand beyond the site of inoculation (Figure 1A). We complemented the *rffG* deletion through *in trans* expression of the *rffG* gene under control of a *lac* promoter for constitutive expression in *P. mirabilis* (23), resulting in the Δ*rffG* p*rffG* strain. The wild-type and the Δ*rffG* strain each carried empty vectors (pBBR1-NheI) to enable growth on the same selective medium as the Δ*rffG* p*rffG* strain. The swarm colonies of the Δ*rffG* p*rffG* strain were attenuated by comparison to the wild-type strain and more expansive than those of the Δ*rffG* strain (Figure 1A), indicating a partial rescue ofswarm motility.

**Figure 1.**
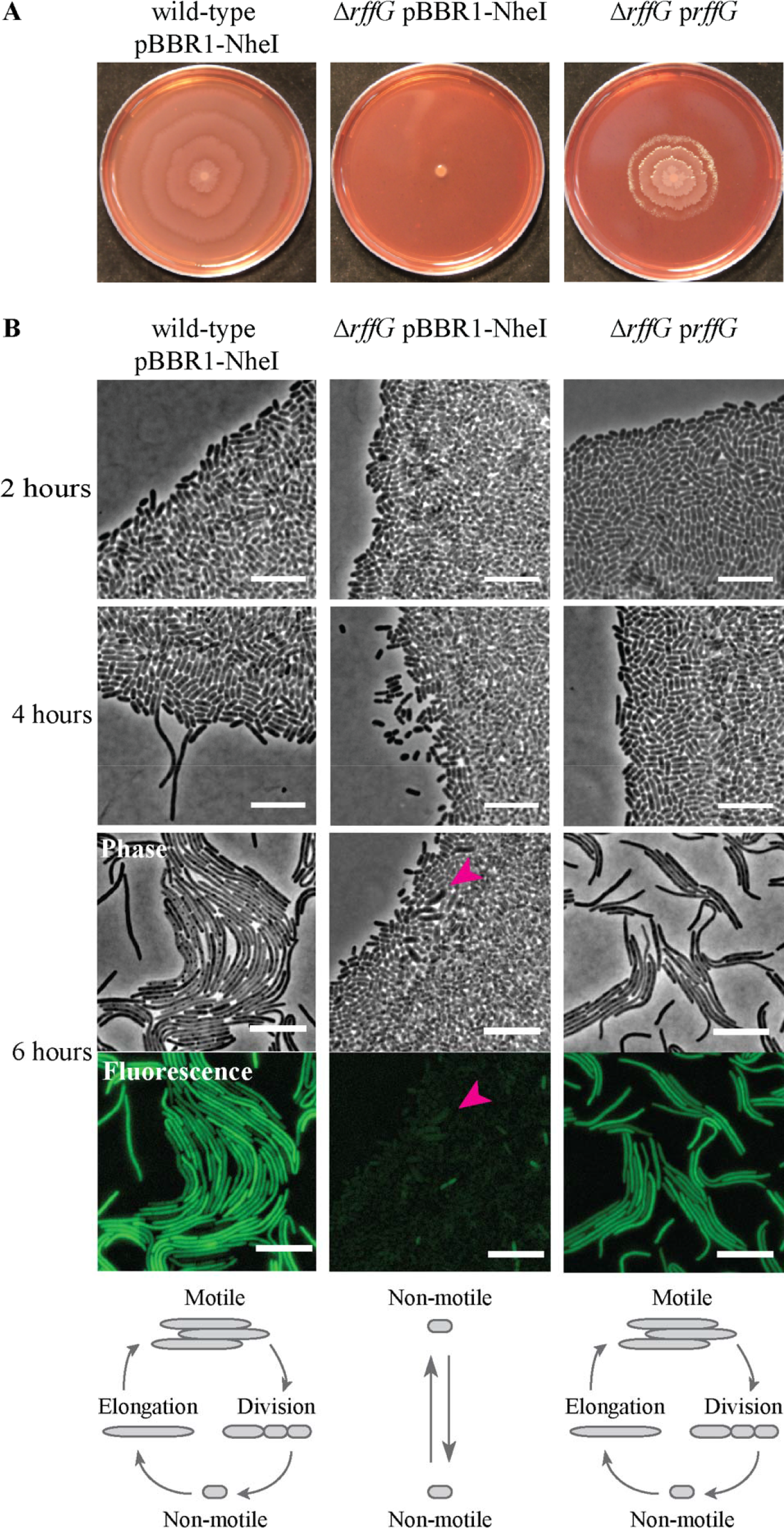
Loss of the *rffG* gene inhibits swarmer cell elongation and swarm motility. A. The wild-type pBBR1-NheI, the Δ*rffG* pBBR1-NheI, and the Δ*rffG* p*rffG* strains were grown on swarm-permissive medium containing kanamycin. B. Phase contrast and epifluorescence microscopy of the BB2000 pBBR1-NheI, the Δ*rffG* pBBR1-NheI, and the Δ*rffG* p*rffG* strains after two, four, and six hours on swarm-permissive medium containing kanamycin. All strains encode Venus immediately downstream of *fliA*. Fluorescence corresponding to *fliA* reporter expression is shown for the six-hour time point. Rolling ball background subtraction was performed using FIJI (72). Magenta arrow highlights an elongating cell in the Δ*rffG* strain that is bulging. Frames from a time-lapse of such cells bulging are in Supplemental Figure SF1E. At bottom are cartoon depictions of the morphological state of cells grown on swarm permissive solid medium. On surfaces, cells elongate up to 40-fold before engaging in motility and dividing into short (1 – 2 µm) non-motile cells. These morphological and behavioral changes coordinate with broad changes to the transcriptome. Scale bars = 10 µm.

We next examined the swim motility of these strains to determine whether loss of *rffG* broadly inhibits flagella-based motility. We analyzed the motility of the wild-type, Δ*rffG*, and Δ*rffG* p*rffG* strains through 0.3% LB agar, which permits swimming. The Δ*rffG* strain, and to a lesser extent the Δ*rffG* p*rffG* strain, was delayed in the initiation of swimming as compared to the wild-type strain. However, all strains occupied the full 10-cm diameter petri dish within 24 hours (Figure SF1A). We measured cell viability in both liquid (Figure SF1B) and in swarms (Figure SF1C) and found that all populations grew to equivalent densities. Thus, the *rffG* gene was essential for surface-based swarm motility, but not for liquid-based swimming motility or growth.

Given that cells required the *rffG* gene to engage in surface-based motility, we hypothesized that cells of the Δ*rffG* strain might fail to progress through stages of swarming such as increased expression of *flhDC*-regulated genes, elongation into swarmer cells, or migration across a surface (Figure 1B). Therefore, we independently assessed the cell morphology of the wild-type, Δ*rffG*, and Δ*rffG* p*rffG* strains using epifluorescence microscopy under swarm-permissive conditions. To visualize flagellar gene expression, a Venus fluorescent protein reporter was introduced on the chromosomes of each strain background downstream of *fliA*. The *fliA* gene encodes the flagellar sigma factor (σ^28^) and is both directly regulated by FlhD_4_C_2_ and highly expressed in swarming cells (36, 37). By four hours after inoculation onto CM55 agar at 37^°^C, populations of the wild-type strain contained many short non-motile and few elongated motile cells (Figure 1B). After six hours, elongated cells expressing the fluorescent *fliA* reporter dominated the inoculum edge and were apparent at the leading edge of the swarms (Figure 1B). By contrast, most cells of the Δ*rffG* strain were short and non-motile at four and six hours; cells appeared modestly shorter than non-elongated wild-type cells (Figure 1B). Cells of the Δ*rffG* strain largely did not exhibit *fliA*-associated fluorescence (Figure 1B). We confirmed the reduction of flagella by visualizing cells harvested from a swarm using transmission electron microscopy. Cells of the Δ*rffG* strain were uniformly short and lacked the extended structures present on the wild-type elongated swarmer cells (Figure SF1D). Notably, some cells of the Δ*rffG* strain did initiate elongation but often ruptured or divided into short cells before completing elongation (Figure SF1E), indicating a failure to complete swarmer cell elongation. By contrast, cells of the Δ*rffG* p*rffG* strain formed elongated motile cells displaying *fliA-*associated fluorescence by six hours (Figure 1B); this progression was delayed as compared to the wild-type strain, which is consistent with a partial rescue. Therefore, as cells of the Δ*rffG* strain failed to increase expression of flagellar genes and to elongate into swarmer cells upon surface contact, we concluded that the *rffG* gene was necessary and sufficient for cells to initiate swarmer cell elongation.

### The *rffG* gene is essential for the production of LPS-associated moieties necessary for cell envelope integrity

We considered that the lack of swarmer cell elongation in the Δ*rffG* strain could be caused by either a physical constraint such as a lack of membrane integrity or by activation of swarm-inhibitory signaling pathways. Therefore, we first examined the membrane composition and integrity of this strain. In *P. mirabilis*, ECA can exists in many forms: linked to other lipids, found in a circularized and soluble form, or surface-exposed and linked to the LPS core in the outer membrane (35, 38-42). Repeated efforts to confirm the presence of ECA via Western blotting with *E. coli* O14 serum (SSI Diagnostica, Hillerød, Denmark), which is reactive against*E. coli*-derived ECA (43, 44), were unsuccessful. We instead characterized overall LPS composition and cell envelope sensitivity to antibiotics. We extracted LPS from surface-grown colonies of the wild-type, Δ*rffG*, and Δ*rffG* p*rffG* strains and then visualized the LPS-associated moieties using silver stain (45). The observed banding patterns of the wild-type and Δ*rffG* p*rffG* strains were nearly equivalent (Figure 2A). The banding pattern of the Δ*rffG* strain, however, lacked a high molecular weight smear, and the bands within the putative O-antigen ladder formed double bands instead of a single band (Figure 2A). We concluded that the *rffG* gene was essential for production of full-length and wild-type LPS, specifically the O-antigen and the high molecular weight components.

We reasoned that these perturbations to the LPS components might cause broader cell envelope damage in cells of the Δ*rffG* strain. To target the outer membrane, we measured resistance to polymyxin B, bile salts, and sodium dodecyl-sulfate (SDS). Polymyxin B is thought to bind LPS, and disruption of LPS biosynthesis genes causes polymyxin B sensitivity in *P. mirabilis* (25, 46). Bile salts (47) and SDS broadly target membranes through detergent-like effects. ECA is likely involved in bile salts resistance of *Salmonella enterica* (48); however, a role for ECA in bile salt resistance of *P. mirabilis* has not been explored. Populations of the wild-type and the Δ*rffG* strains were resistant to fully saturated solutions (50 mg/mL) of polymyxin B (Table 1; Figure SF2A) and exhibited reduced growth on 0.5% bile salts (Table 1; Figure SF2B). However, the growth defects of the Δ*rffG* strain on 0.2% bile salts were more severe than those of the wild-type strain. The Δ*rffG* strain was reduced in growth and formed small and translucent colonies (Figure SF2B). The Δ*rffG* strain was also more sensitive to SDS than the wild-type strain. 0.5% SDS was permissive for growth of the wild-type strain but not for the Δ*rffG* strain (Table 1; Figure SF2C). As controls, we measured sensitivity to the non-membrane targeting antibiotics, gentamycin and kanamycin. We found no differences in growth between the wild-type and the Δ*rffG* strains when grown on gentamycin and kanamycin (Table 1). In sum, the *rffG*-deficient cells exhibited increased sensitivity to bile salts and SDS, but not to polymyxin B, gentamycin, and kanamycin. Therefore, the outer membrane in *rffG*-deficient cells was compromised in a phenotypically distinct manner than previously studied LPS-deficient *P. mirabilis* strains.

**Figure 2.**
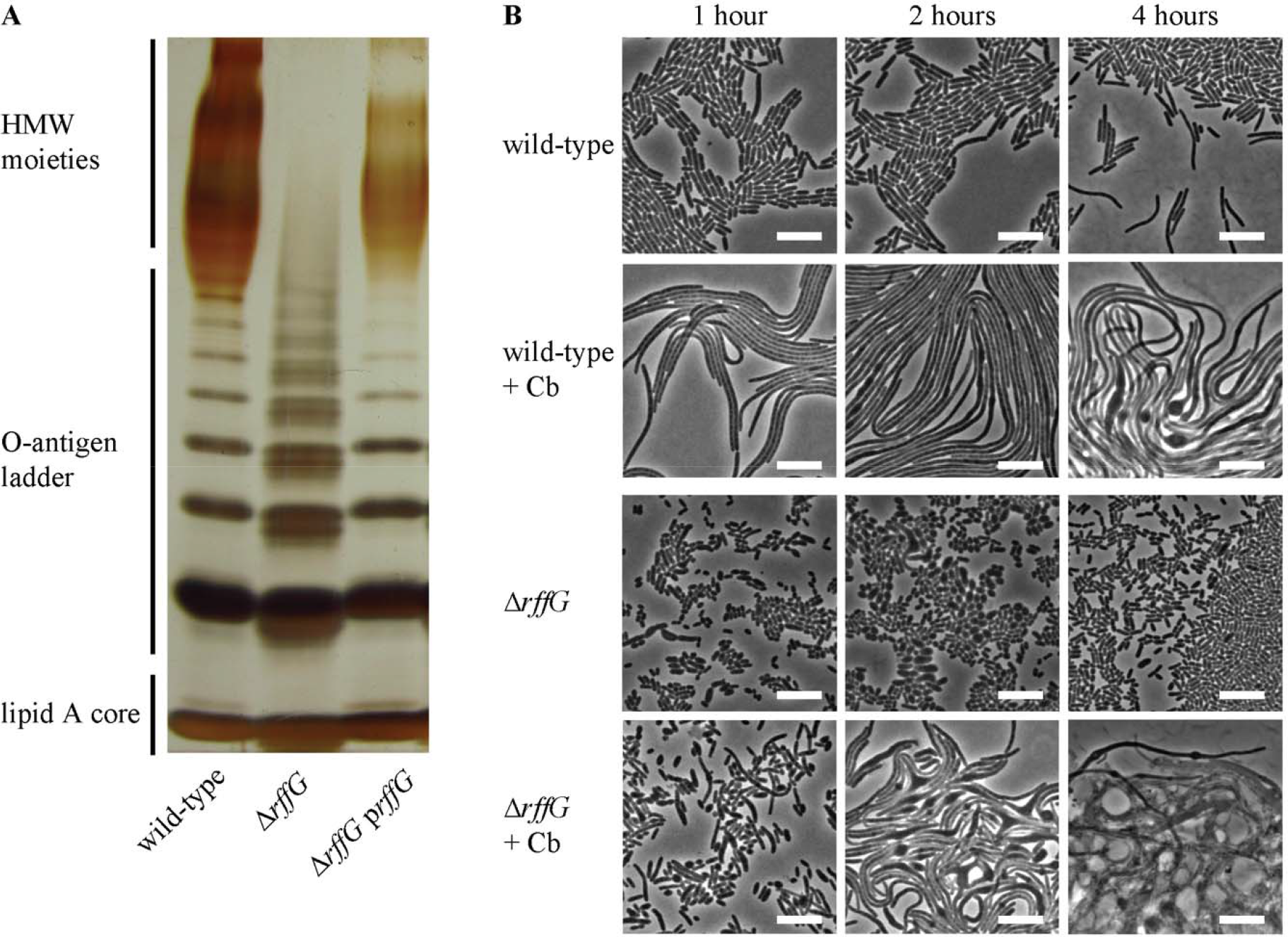
Loss of the *rffG* gene affects outer membrane structures and cell envelope integrity. A. LPS was extracted from surface-grown cells of the wild-type, the Δ*rffG*, and the Δ*rffG* p*rffG* strains. Samples were run on a 12% SDS-PAGE gel and detected via silver stain. Predicted LPS-associated moieties are labeled on the left. HMW = high molecular weight. B. The wild-type and the Δ*rffG* strains were spread onto swarm-permissive agar pads containing 0 or 10 µg/mL carbenicillin (Cb). Shown are cells after one, two, or four hours of incubation on a surface. Images are representative; N = 3. Scale bars = 10 µm.

**Table 1:** Antibiotic sensitivity of the wild-type and the Δ*rffG* strains on surfaces.

| | Polymyxin B<br>MIC, mg/mL | Bile salt<br>MIC, % w/v | SDS<br>MIC, % w/v | Gentamycin,<br>MIC, μg/mL | Kanamycin<br>MIC, μg/mL |
| --- | --- | --- | --- | --- | --- |
| the wild-type<br>strain | > 50 | 0.2 | 0.05 | 0.1 | 0.1 |
| the $\Delta rffG$ strain | > 50 | 0.1 | 0.01 | 0.1 | 0.1 |

To further interrogate cell envelope integrity, we analyzed cell morphology in response to a subinhibitory concentration (10 µg/ml) of the beta-lactam antibiotic carbenicillin. Within one hour of growth on carbenicillin-containing CM55 agar at 37°C, we observed that wild-type cells lengthened to tens of microns while remaining a uniform diameter (Figure 2B). By four hours of growth, occasional bloating was apparent at the mid-cell (Figure 2B). By contrast after one hour of growth on carbenicillin-containing CM55 agar at 37°C, many cells of the Δ*rffG* strains remained short; the few elongated cells appeared wider or lemon-shaped (Figure 2B). After two hours of growth, the population of the Δ*rffG* strain consisted of elongated, bloated cells, including several with triangular protrusions (Figure 2B). By four hours of growth, most cells of the Δ*rffG* strain had ruptured; the remaining cells were several microns long (Figure 2B). Equivalent results were attained with Aztreonam, an inhibitor of the cell division protein, FtsI (Figure SF2D). The *rffG*-deficient cells were therefore more susceptible to cell wall stress and membrane-targeting detergents, indicating that the composition of the outer membrane in *rffG-*deficient cells was compromised.

### The *rffG* deficiency induces stress responsive pathways in cells

Loss of the *rffG* gene resulted in altered cell envelope structure (Figure 2A) and stability (Figures 2B, SF2D), both of which would likely induce stress-responsive pathways. Interestingly, the *rffG*-deficient cells could transiently elongate when artificially driven to expand in length using carbenicillin; therefore, inhibited elongation in these cells was not purely due to disrupted physical structures of the cell envelope. Such cell envelope stress in *P. mirabilis* can activate several swarm-inhibitory pathways such as those controlled by RcsB, RpoE, and RppA/B (22, 24, 25, 30). To identify whether one or more of these pathways was induced in *rffG*-deficient cells, we performed RNA-Seq analysis on cells of the wild-type and Δ*rffG* strains harvested under swarm-permissive conditions. Two biological replicates were acquired and examined. Short, non-motile cells harvested from swarms of the wild-type strain were used as the control for these experiments.

343 genes out of ~3455 protein-coding genes in the *P. mirabilis* BB2000 genome were expressed at least four-fold differently (Figure SF3). 136 genes were decreased in the Δ*rffG* strain, of which approximately 18% of were related to ribosome structure and translation, and 20% were directly related to flagella assembly or chemotaxis (Table S1). Additional factors known to regulate swarmer cell development and motility also had decreased expression, including *umoD* at 0.12-fold, *umoA* at 0.19-fold, and *ccm* at 0.12-fold (Table 2). Cell-envelope associated genes were also down-regulated, e.g., the penicillin-binding protein gene *pbpC* and the membrane lipid modifying gene *ddg* at 0.25 and 0.22-fold, respectively (Table S1). By contrast, 207 genes were increased, including the virulence-associated MR/P fimbria (200-fold increased expression of *mrpA*) and *Proteus* P-like pili (55.5-fold increased expression of *pmpA*) (Tables 3 and S2). In addition, several genes related to carnitine metabolism were increased, including the transcriptional regulator *caiF* at 63.6-fold and *caiA, fixC*, and *fixX* at four-fold (Tables 3 and S2). Carnitine can be metabolized, particularly under anaerobic conditions (49-51), and can act as a stress protectant for several bacterial species (52, 53) (also reviewed in (54)). Likewise, *ompW* was increased ~ 33.7 fold along with *dcuB*, an anaerobic C4-dicarboxylate transporter, at 29.6 fold (Tables 3 and S2). In *E. coli*, maximal *ompW* expression is tied to survival in the transition from aerobic to anaerobic growth (55). Therefore, genes for fimbrial production and for metabolism under anaerobic or micro-aerobic environments were more highly expressed in the *rffG*-deficient cells; by contrast, genes promoting swarm motility were decreased.

Notably, three major regulators were expressed much higher in the Δ*rffG* strain (Table 2): *mrpJ* at 19.1-fold, *rpoS* at 8.7-fold, and the RcsB-cofactor *rcsA* at 9.9-fold. Both *mrpJ* and *rcsA* were previously shown to contribute to swarm inhibition (6, 7, 34). We observed a partial overlap between differentially regulated genes and the characterized MrpJ regulon (34). However, we found a larger overlap between the genes differentially regulated in the Δ*rffG* populations with the genes recently characterized as regulated by RcsB in *P. mirabilis* (Tables 2 and 3) (6, 7). The overlap was especially striking among down-regulated genes. While about 8% of up-regulated genes overlapped with the Rcs regulon, 24% of down-regulated genes overlapped with the Rcs regulon (Figure SF3). RcsB directly represses *flhDC*, which in turn regulates flagella and chemotaxis genes, positively regulates paralogues of the swarm-inhibitory *mrpJ* gene (6, 7), and regulates *minCDE* (6). Therefore, we hypothesized that the Rcs phosphorelay was likely activated in the *rffG*-deficient cells.

**Table 2:** Top 20 genes down-regulated in the Δ*rffG* strain relative to the wild-type strain.

| Transcript | Product | Fold<br>Change | Rcs<br>Regulon* |
| --- | --- | --- | --- |
| *cspB* | cold shock protein | 0.01 |  |
| *flgN* | flagella synthesis protein | 0.03 |  |
| *BB2000_1016*,<br>*BB2000_1017* | cold shock protein, heat shock protein | 0.04 |  |
| *flgM* | anti-sigma28 factor FlgM | 0.04 |  |
| *BB2000_3499* | lipoprotein | 0.05 | Yes |
| *fliE* | flagellar hook-basal body complex<br>protein | 0.05 |  |
| *BB2000_1271* | putative MFS-type transporter YdeE | 0.07 |  |
| *fliA* | flagellar biosynthesis sigma factor | 0.08 | Yes |
| *BB2000_1466* | hypothetical protein |  | Yes |
| *holD* | DNA polymerase III subunit psi | 0.10 | Yes |
| *intB* | prophage integrase | 0.11 |  |
| *fliM, fliN* | flagellar motor switch protein FliM,<br>flagellar motor switch protein FliN | 0.11 | Yes |
| *BB2000_0342* | transcriptional regulator | 0.11 | Yes |
| *BB2000_0996,*<br>*umoD* | hypothetical protein, exported protein<br>(upregulator of flagellar master operon) | 0.12 | Yes |
| *rpsO* | 30S ribosomal protein S15 | 0.12 |  |
| *ccm* | membrane protein (Ccm1 protein) | 0.12 | Yes |
| *rplI* | 50S ribosomal protein L9 | 0.12 |  |
| *fliZ* | flagella biosynthesis protein FliZ | 0.12 | Yes |
| *BB2000_2557* | phospholipid-binding protein | 0.13 | Yes |
| *BB2000_2289,*<br>*fruK* | hypothetical protein, 1-phosphofructokinase | 0.13 |  |

\* Based on the published RcsB regulon in *P. mirabilis* (6, 7)

**Table 3:** Top 20 genes up-regulated in the Δ*rffG* strain relative to the wild-type strain.

| **Transcript** | **Product** | **Fold**<br>**Change** | **Rcs**<br>**Regulon\*** |
| --- | --- | --- | --- |
| *mrpA* | major mannose-resistant/Proteus-like<br>fimbrial protein | 200.27 | Yes |
| *caiF* | DNA-binding transcriptional activator<br>CaiF | 63.60 |  |
| *pmpA* | fimbrial subunit | 55.46 |  |
| *BB2000_1924* | transcriptional regulator | 45.63 |  |
| *BB2000_1499* | fimbrial subunit | 44.85 |  |
| *BB2000_3017* | fimbrial operon regulator | 44.59 |  |
| *zntB* | zinc transporter | 42.83 |  |
| *mrpG* | fimbrial subunit | 42.72 |  |
| *BB2000_0667* | hypothetical protein |  |  |
| *ompW* | outer membrane protein W | 33.70 |  |
| *BB2000_2725* | TetR-family transcriptional regulator | 32.64 |  |
| *dcuB* | anaerobic C4-dicarboxylate transporter | 29.58 |  |
| *BB2000_0531* | sigma 54 modulation protein | 25.53 |  |
| *BB2000_0331* | heat shock protein HtpX | 22.81 |  |
| *BB2000_0299* | hypothetical protein | 22.47 |  |
| *BB2000_0395* | hypothetical protein | 21.42 | Yes |
| *BB2000_0229* | hypothetical protein | 21.19 |  |
| *BB2000_1497* | fimbrial chaperone protein | 20.97 |  |
| *mrpJ* | fimbrial operon regulator | 19.08 |  |
| *dmsA* | dimethyl sulfoxide reductase chain A | 18.99 | Yes |

\* Based on the published RcsB regulon in *P. mirabilis* (6, 7)

### RcsB inhibits swarmer cell elongation and morphology defects of *rffG*-deficient cells

We hypothesized that RcsB-mediated repression of the swarm transcriptional regulator *flhDC* was the primary cause for loss of swarmer cell elongation and swarm motility of the Δ*rffG* strain. Therefore, we independently constructed a chromosomal deletion of *rcsB* in the wild-type and the Δ*rffG* strains, resulting in the Δ*rcsB* and Δ*rffG*Δ*rcsB* strains, respectively. We also constructed a plasmid for constitutive and increased expression of *flhDC* and introduced this *in trans* in the wild-type and the Δ*rffG* strains, resulting in BB2000 p*flhDC* and the Δ*rffG* p*flhDC* strain, respectively. As RcsB is an upstream repressor of *flhDC*, we predicted that swarm motility and swarmer cell elongation would be rescued in both the Δ*rffG*Δ*rcsB* and the Δ*rffG* p*flhDC* strains. The Δ*rcsB* and BB2000 p*flhDC* strains were predicted to contain constitutively swarming cells based on equivalent constructs in other *P. mirabilis* wild-type backgrounds (9, 29). We inoculated all strains onto separate swarm-permissive agar plates and analyzed colony expansion over 24 hours growth at 37^°^C (Figure 3). As predicted, the Δ*rcsB* and BB2000 p*flhDC* strains expanded across the 10-cm diameter plate by 16 hours (Figure 3A). However, neither the Δ*rffG*Δ*rcsB* nor Δ*rffG* p*flhDC* strains expanded beyond 20% of the plate by 16 hours (Figure 3A). The Δ*rffG* p*flhDC* strain did reach the edge of plate by 24 hours, but the Δ*rffG*Δ*rcsB* strain remained constrained towards the center (Figure 3B). Extracted LPS of these strains grown in liquid broth and on surfaces were analyzed. We found that the banding pattern of the Δ*rcsB* strain was equivalent to that of the wild-type strain, BB2000 (Figure SF4). Likewise, the banding pattern of the Δ*rffG*Δ*rcsB* and the Δ*rffG* p*flhDC* strains were equivalent to that of the Δ*rffG* strain (Figure SF4). Thus, neither RcsB nor FlhD_4_C_2_ contributed to the production of the *rffG-*dependent LPS-associated moieties. However, increased expression of *flhDC* or deletion of *rcsB* was sufficient to increase swarm motility of *rffG*-deficient cells, indicating that RcsB-mediated repression of *flhDC* was sufficient for the swarm inhibition of *rffG*-deficient cells.

**Figure 3.**
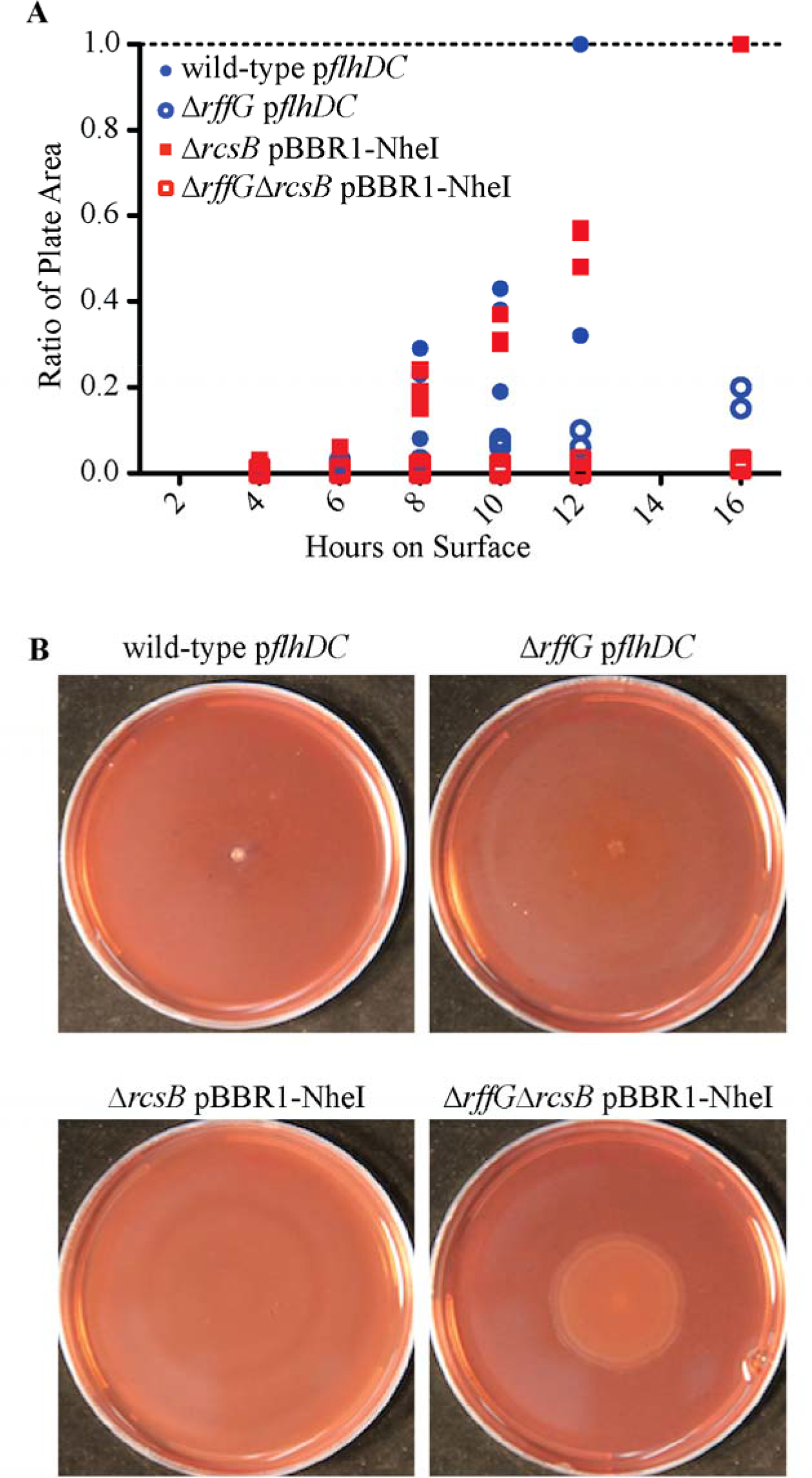
Loss of the *rffG* gene impacts swarm colony development. A. Loss of the *rffG* gene extends the lag phase before swarm colony expansion. Liquid-grown populations of the wild-type p*flhDC*, the Δ*rffG* p*flhDC*, the Δ*rcsB* pBBR1-NheI, and the Δ*rffG*Δ*rcsB* pBBR1-NheI strains were normalized based on OD_600_ and inoculated onto swarm permissive plates containing kanamycin for plasmid retention. Area of visible swarm colony expansion was measured at indicated times; each time-point comprised of separate plates. N = 3 for each strain and time-point. B. Populations of the wild-type p*flhDC*, the Δ*rffG* p*flhDC*, the Δ*rcsB* pBBR1-NheI, and the Δ*rffG*Δ*rcsB* pBBR1-NheI strains were grown on a swarm-permissive plates containing kanamycin for plasmid retention. Populations of the wild-type p*flhDC* (N = 3), the Δ*rffG* p*flhDC* (N = 8), and the Δ*rcsB* pBBR1-NheI (N = 4) strains form a thin film with no apparent concentric rings. Populations of the Δ*rffG*Δ*rcsB* pBBR1-NheI strain (N = 4) produced mucoid swarm colonies that formed tight concentric rings. Representative images are shown.

Since the Δ*rffG*Δ*rcsB* strain did not have full recovery of swarm motility by 24 hours, we hypothesized that the loss of the *rcsB* gene might affect a pathway separate from the FlhD_4_C_2-_regulated genes. We integrated the chromosomal *fliA*-Venus transcriptional reporter into each strain and observed the resultant cells using epifluorescence microscopy under swarm-permissive conditions. After six hours of growth at 37°C, the BB2000 p*flhDC* -derived and the Δ*rffG* p*flhDC*-derived strains consisted of motile, elongated cells with *fliA* reporter-associated fluorescence (Figure 4). Likewise, cells in the Δ*rcsB* strain were generally elongated and motile with *fliA* reporter-associated fluorescence (Figure 4). Surprisingly, cells of the Δ*rffG*Δ*rcsB* strain exhibited severe cell shape defects: cells were bloated and uneven in width, forming spheres, tapering at the cell poles, or bulging at the mid-cell (Figure 4). In addition, several cells of the Δ*rffG*Δ*rcsB* strain were forked at the cell pole or branched at the mid-cell, resulting in the formation of more than two cell poles. Nonetheless, the elongated cells of the Δ*rffG*Δ*rcsB* strain exhibited *fliA* reporter-associated fluorescence and were motile (Figure 4). In sum, cells of the Δ*rffG*Δ*rcsB* strain did not retain fidelity of a two-pole, rod-shaped swarmer cell morphology, even though they had increased *fliA* expression. Thus, RcsB contributed to the suppression of shape defects in *rffG*-deficient cells. As these defects only arose in the absence of RcsB, we posit this was achieved via RcsB-dependent and FlhD_4_C_2_*-*independent pathway(s).

**Figure 4.**
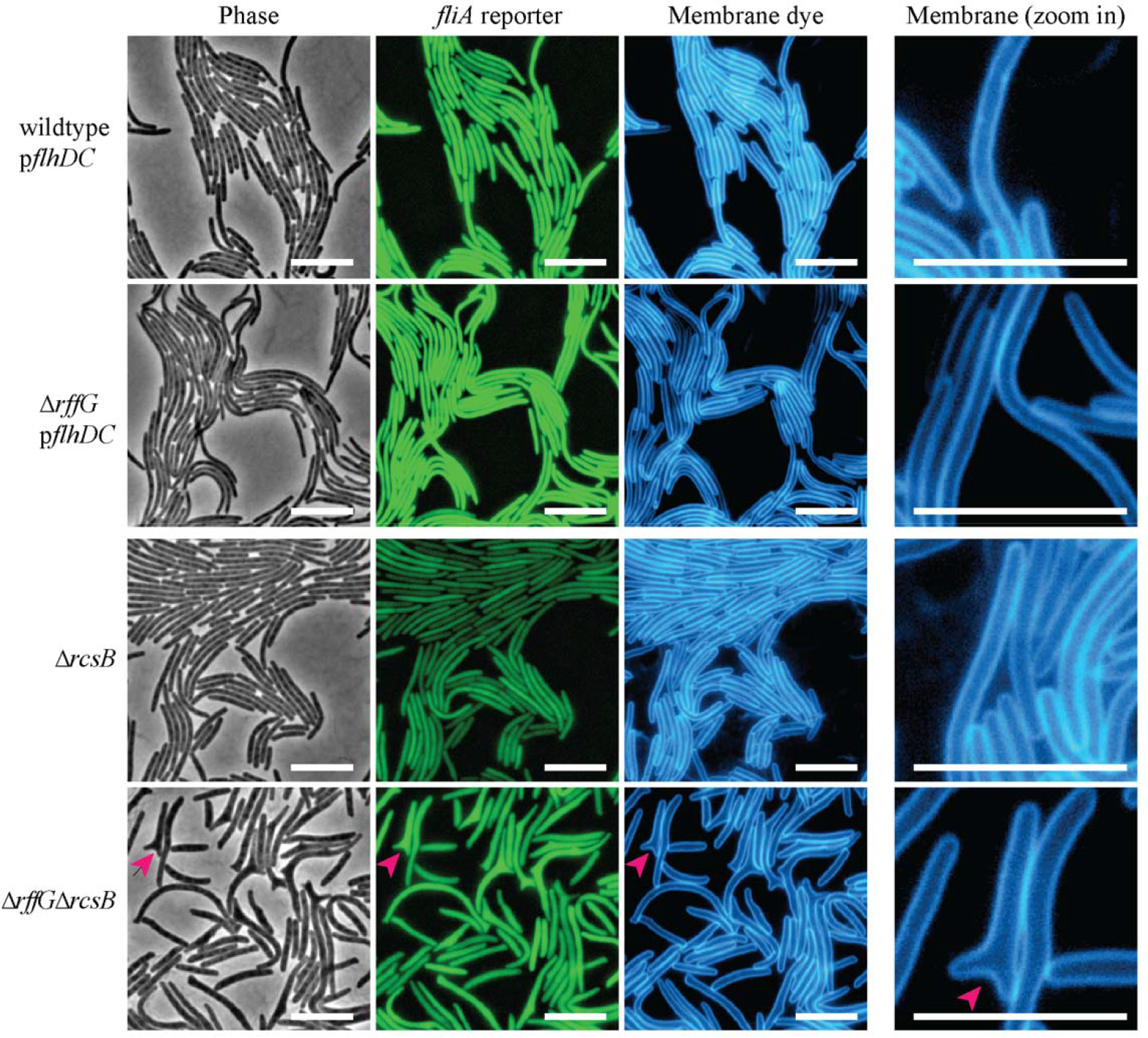
Swarm cell elongation of Δ*rffG* cells are rescued by increased *flhDC* expression or deletion of the *rcsB* gene. Epifluorescence microscopy of the wild-type p*flhDC*, the Δ*rffG* p*flhDC*, the Δ*rcsB*, and the Δ*rffG*Δ*rcsB* strains on swarm permissive agar pads. All strains encode Venus controlled by the promoter for *fliA*. For populations of the wild-type p*flhDC* and the Δ*rffG* p*flhDC* strains, the agar contained kanamycin for plasmid retention. Images were taken once swarm motility initiated (four to six hours on surface). Rolling ball background subtraction on *fliA* reporter images was conducted using FIJI (63). Magenta arrows indicate a cell exhibiting shape and polarity defects. Cropped images are shown on the far right for emphasis on cell shape defects. Left, phase contrast; middle left, Venus expression; middle right, membrane stain; far right, cropped selection of membrane stain image at higher zoom. Scale bars = 10 µm.

## Discussion

Here crucial insights were elucidated about the role of cell envelope structure and stress sensing in the development of *P. mirabilis* swarmer cells, specifically regarding the cell envelope biosynthesis gene *rffG* and the signaling pathways that respond to its absence (Figure 5). We have shown that the *rffG* gene was essential for the assembly of a swarm-permissive cell envelope. Loss of the *rffG* gene resulted in the loss of LPS-associated moieties, the alteration of the O-antigen ladder, and increased sensitivity to antimicrobials that specifically target the cell envelope. Based on the RNA-Seq results, cells of the Δ*rffG* strain entered into a distinctive transcriptional state, resulting in the upregulation of several stress responsive pathways. While most of the pathways activated by loss of *rffG* have yet to be characterized, RcsB-mediated inhibition of *flhDC* expression was a major regulatory factor in restricting elongation in the Δ*rffG* strain. Loss of *rcsB* or over-expression of *flhDC* rescued swarm motility in Δ*rffG* cells. Moreover, an additional role for RcsB in the maintenance of cell shape and polarity during swarmer cell elongation was uncovered as RcsB served to also maintain the two-pole, rod shape of *rffG*-deficient cells.

**Figure 5.**
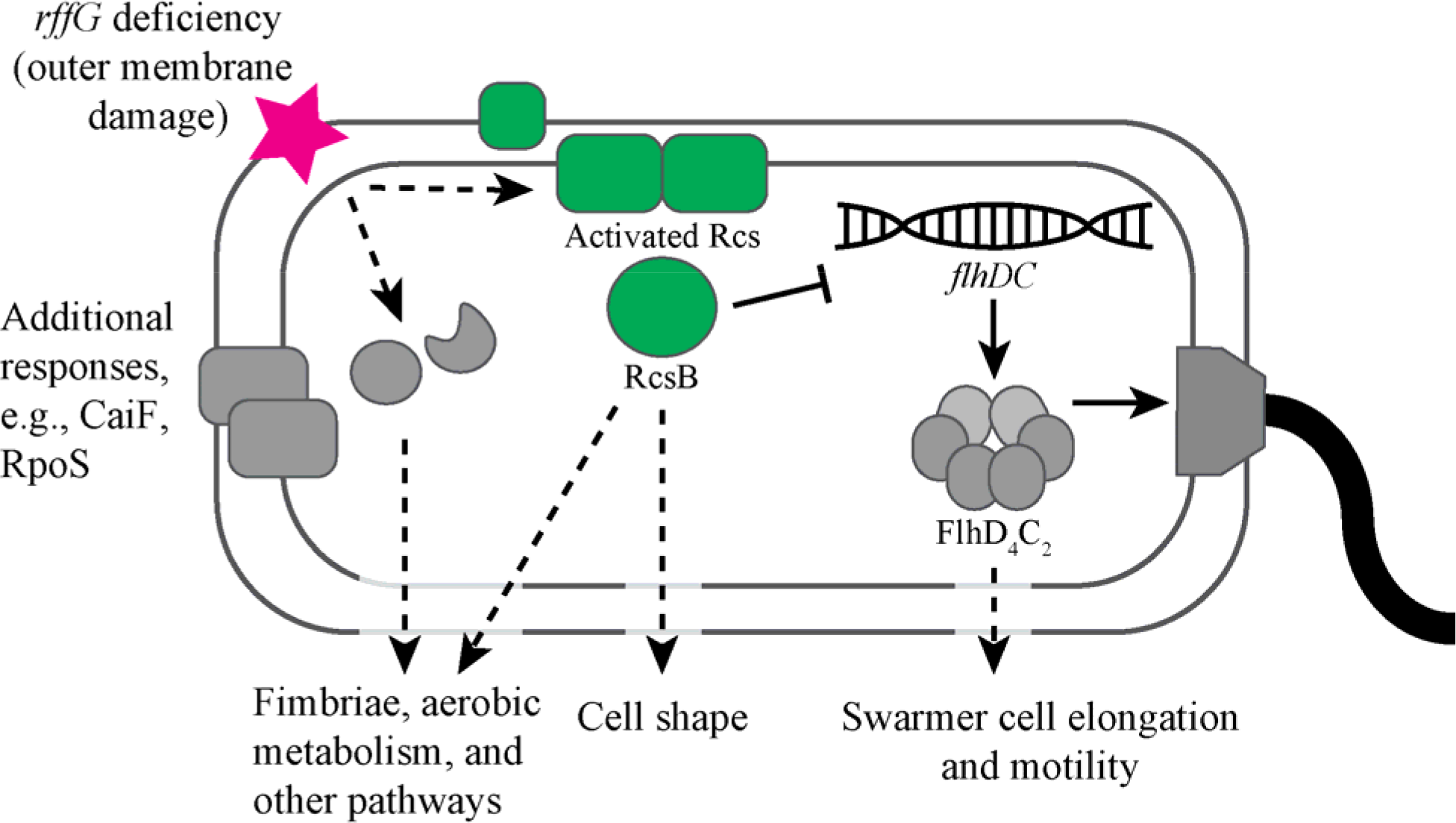
Checkpoint model in which a *rffG* deficiency induces multiple stress-associated pathways controlling swarmer cell elongation and cell shape. Loss of the *rffG* gene induced activity of RcsB which in turn directly represses expression of the *flhDC* genes that encode the master regulator of flagella production and swarm motility. We found that RcsB in *P. mirabilis* was also necessary to maintain cell shape and polarity through yet uncharacterized mechanisms, supporting a role for RcsB as a multi-functional regulator of swarmer cell development and motility. The composition of the cell envelope, including the *rffG*-dependent moieties, may serve as a developmental checkpoint before engaging in swarmer cell development. Cell elongation and increased *flhDC* expression are initial steps in swarmer cell development.

The transcriptional state of cells in the Δ*rffG* strain was characterized by the activation of pathways such as those controlled by the transcriptional regulators RpoS, CaiF, and RcsB. There was increased expression of several fimbrial gene clusters. There was also a notable increase in carnitine metabolism genes, which is associated with growth under anaerobic conditions (50, 51). Many of the identified genes are controlled by MrpJ and oxygen availability (56), raising the possibility that cells of the Δ*rffG* strain might bias towards a more adherent, low-oxygen lifestyle. Flagellar and chemotaxis genes had decreased expression in the Δ*rffG* strain. The disruption of the flagellar pathway was consistent with the loss of swarm motility in the Δ*rffG* strain. However, potential mechanisms for inhibiting cell elongation and driving cell shape defects were less apparent in the RNA-Seq data. For example, while RcsB has been implicated in cell elongation via regulation of *minCDE* (6, 7), differential regulation of these genes in the Δ*rffG* strain was not evident. Further research will need to be done to completely categorize the genes differentially regulated in *rffG*-deficient cells and to fully understand the physiological and behavioral implications of these altered expression levels.

Several questions remain regarding the mechanisms of activation, as well as the downstream activity, of RcsB and MrpJ in Δ*rffG* cells. First, Bode et al recently demonstrated that MrpJ acts as a regulator mediating the transition of cells between swarm motility (MrpJ-repressed) and non-motile adherence (MrpJ-induced) similar to RcsB (34). MrpJ and RcsB may positively regulate each other and have overlapping regulons (6, 7, 34), making it difficult to genetically disentangle the contributions of each regulator. Additionally, perturbation of outer membrane structures appears to be communicated to the Rcs phosphorelay through both RcsF-dependent and independent pathways in *P. mirabilis* (22). Whether cell envelope stress of *rffG-*deficient cells is communicated through the outer membrane-localized RcsF, through the upregulation of RcsA, or through an uncharacterized additional pathway remains to be determined. Also unknown is whether the *rffG-*dependent LPS-associated moieties communicate to Rcs via the Umo system as was previously shown for O-antigen (22).

Previous research has elucidated how disrupting LPS induces stress-responsive pathways leads to swarm inhibition. For example, abrogation of O-antigen structure through deletion of the O-antigen ligase (*waaL*) or chain length determinant (*wzz*) inhibits activation of *flhDC* upon surface contact (23). And loss of the sugar-modifying O-antigen biosynthesis genes *ugd* and *galU* inhibits swarmer cell elongation and motility (25). The aforementioned genes are implicated in LPS biosynthesis. Here we propose that *P. mirabilis* cells require cell envelope structures in addition to LPS for the initiation of swarmer cell elongation (Figure 5). The *rffG-*dependent high molecular weight LPS-associated moieties are not chemically characterized; we hypothesize that these might consist of LPS-associated ECA or ECA-derived moieties since the *E. coli rffG* homologue is needed for the production of the broadly conserved ECA (35). Further research is needed to characterize the structural changes to the outer membrane in *rffG*-deficient cells, especially as these moieties contribute to overall membrane integrity on surfaces.

Outer membrane structure also plays a crucial mechanical role in resisting turgor pressure fluctuations associated with cell wall stress, specifically beta-lactam drugs (57). Cells of the Δ*rffG* strain were sensitive to detergent-like membrane-targeting antimicrobials, altogether suggesting that the *rffG-*dependent moieties are crucial for outer membrane composition and integrity. One explanation is that these *rffG-*associated cell envelope defects are caused by pleiotropic effects resulting from disrupting a cell envelope biosynthesis pathway that uses a shared pool of precursor molecules. We raise this possibility because perturbation of ECA or LPS biosynthesis genes in *E. coli* cause the accumulation of dead-end intermediates that broadly impact cell envelope integrity (20, 21). However, we instead posit that the cell envelope defects in *rffG-*deficient cells might be sufficient to sensitize cells to form defective cell shapes.

We propose that RcsB acts to suppress cell wall defects in *rffG*-deficient cells as well as potentially other cell-envelope compromised (Figure 5). We observed that absence of the *rffG-*dependent moieties did not mechanically restrict swarmer cell elongation or result in the formation of over two poles in artificially elongated cells constitutively expressing *flhDC*. Cells lacking both *rffG* and *rcsB* cells, however, exhibited growth from the mid-cell and the formation of over two cell poles in addition to other physical perturbations. Thus, though deletion of *rcsB* rescues cell elongation and motility through de-repression of *flhDC*, the absence of RcsB also perturbed a yet uncharacterized morphology-generating pathway critical for the cell shape and integrity of *rffG*-deficient cells. Others have also proposed that the Rcs phosphorelay plays a conditional role in cell shape maintenance in other bacteria. L-form *E. coli* cells require the Rcs phosphorelay to recover a rod shape; cells lacking this response rupture (58). The authors of that study proposed that Rcs might function to maintain cell shape in conditions in which cells lose cell wall through exposure to lysozyme or cationic antimicrobial peptides, including several niches within a human host (58). Moreover, in *E. coli* and *Agrobacterium tumerfaciens*, cell polarity defects, which are similar to those of the Δ*rffG*Δ*rcsB* strain, appeared to arise from the formation of patches of inert peptidoglycan and mislocalized division planes (59-63). Further study is needed to mechanistically understand which aspects of the RcsB regulon are specific for cell shape maintenance and how additional poles emerge in cells lacking both RcsB and the *rffG-*dependent moieties.

It remains unclear how RcsB, which is presumably inactive in swarming cells, can play a role in swarmer cell shape maintenance. We propose two broad mechanisms that may resolve this contradiction. First, the role(s) of RcsB in cell shape maintenance may occur prior to initiation of swarmer cell elongation. Elongation may exacerbate unrepaired envelope flaws that manifest in cell shape and polarity defects. As such, the Rcs phosphorelay would act as a developmental checkpoint to restrict swarmer cell development in conditions challenging to the cell envelope. Second, RcsB may have multiple states beyond simply “active” and “inactive” that may allow differential activity across time and cell states. The DNA-binding activity of RcsB has been shown to be modulated by both phosphorylation and association with auxiliary transcription factors in *E. coli* (64). How RcsB activity is modulated downstream of phosphorylation and association with potential auxiliary transcription factors remains unknown in *P. mirabilis*.

Altogether, we propose that cell envelope stress, including the presence of *rffG-*dependent moieties, functions as a developmental checkpoint before swarmer cell elongation and increased flagellar gene expression (Figure 5). Under swarm-permissive conditions in the presence of wild-type cell envelope structure, the Rcs phosphorelay, along with other stress-sensing pathways, would be inactive thereby allowing the swarmer development to progress. When the cell envelope is perturbed, we posit that the activity of cell envelope stress-responsive sensors culminates in the adaptation of an adherence-promoting lifestyle that may provide protection against external stressors. As the Rcs phosphorelay, *flhDC*, and *rffG*, among other discussed genes, are conserved among the Enterobacteriaceae family, we predict these factors may broadly serve to modulate bacterial swarm motility and potentially cell development.

## Experimental Procedures

### Growth conditions

Liquid cultures were grown in LB-Lennox broth. Colonies were grown in 0.3 % LB agar for swimming motility assays, on LSW-agar (65) for plating non-motile colonies, and on CM55 blood agar base (Oxoid, Hampshire, UK) for swarming. Antibiotics for selection were used throughout all assays as following: 15 µg/mL tetracycline (Amresco Biochemicals, Solon, OH), 25 µg/mL streptomycin (Sigma Aldrich, St Louis, MO), and 35 µg/mL kanamycin (Corning, Corning, NY). For swarm assays, overnight cultures were normalized to OD_600_ 1.0, and 1 µl of culture was inoculated with a needle onto swarm-permissive CM55 blood agar base (Oxoid, Hampshire, UK) plates containing 40 µg/mL Congo Red, 20 µg/mL Coomassie Blue, and kanamycin (Corning, Corning, NY) as needed. Plates were incubated at 37°C. When indicated, we used strains carrying an empty vector (pBBR1-NheI (66)) to confer kanamycin resistance. Images were taken with a Canon EOS 60D camera.

### Strain construction

Strain construction was performed as described previously (67). Strains and plasmids are listed in Table 4. All plasmids were confirmed by Sanger Sequencing (Genewiz, South Plainfield, NJ). For all strains, expression plasmids were introduced into *P. mirabilis* via *E. coli* SM10λpir as previously described (65). Resultant strains were confirmed by Polymerase Chain Reaction (PCR) of the targeted region. The Δ*rffG* strain was additionally confirmed through whole genome sequencing as described in (68).

**Table 4:**
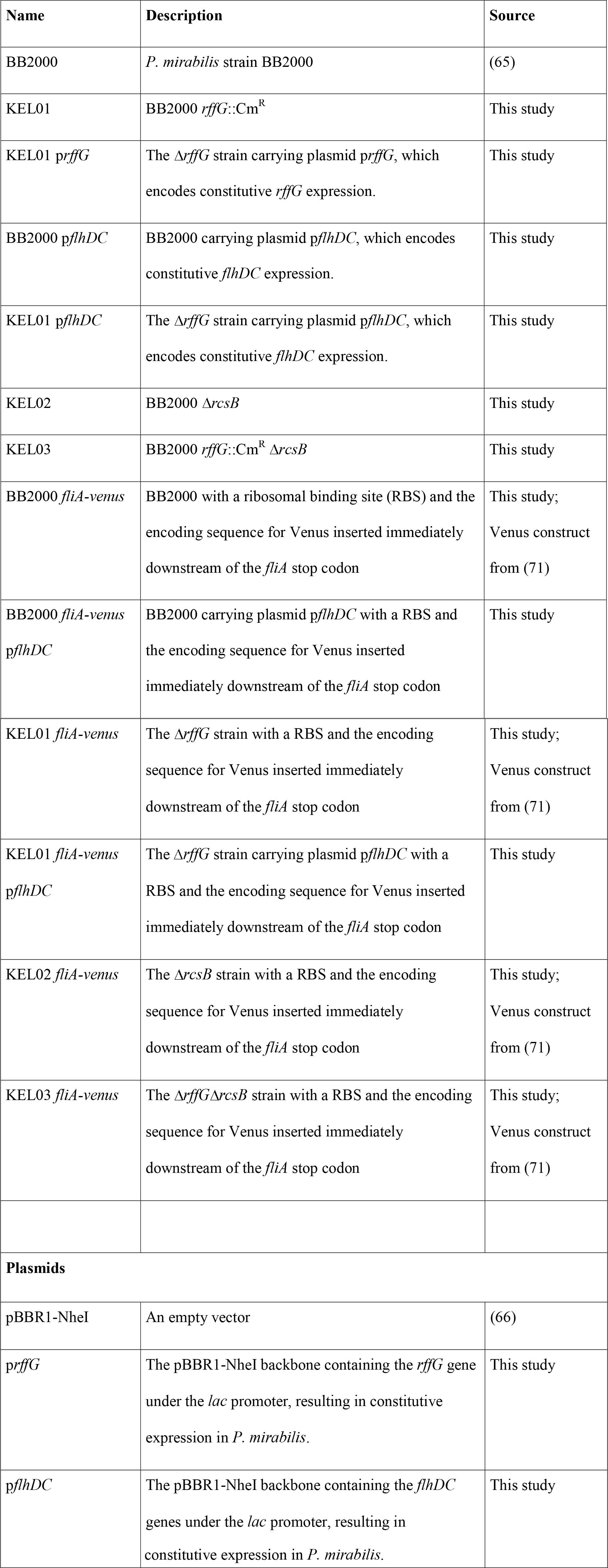
Strains and plasmids used in this study.

For construction of the Δ*rcsB* strain, a gBlock (Integrated DNA Technologies, Coralville, IA) containing the 452 base-pairs (bp) upstream and downstream of *rcsB* (*P. mirabilis* BB2000, accession number CP004022:nt 1972701…1973153 and 1973809…1974261) was generated and introduced to pKNG101 at SpeI and XmaI sites using SLiCE (69). Similarly, for construction of the Δ*rffG* strain, a gBlock containing a chloramphenicol resistance cassette (amplified from pBAD33 (70)) flanked by 1000 bp upstream of *rffG* and downstream of *rffG* (*P. mirabilis* BB2000, accession number CP004022:nt 3635400…3636399 and 3637473…3638473) (Integrated DNA Technologies, Coralville, IA) was introduced to pKNG101 (67) at the same sites. For construction of *fliA* reporter strains, a gBlock encoding the last 500 bp *fliA* (*P. mirabilis* BB2000, accession number CP004022:nt 1856328…1856828), RBS (aggagg), a modified variant of Venus fluorescent protein (a gift from Drs. Enrique Balleza and Philippe Cluzel (71), and 500bp downstream *fliA* (*P. mirabilis* BB2000, accession number CP004022:nt 1855828…1856328) (Integrated DNA Technologies, Coralville, IA) was inserted into pKNG101 (67) at the ApaI and XbaI sites. For construction of the *rffG* expression strains, the nucleotide sequence for *rffG* was amplified via PCR from the *P. mirabilis* chromosome using oKL273 and oKL274 and inserted into expression vector pBBR1-NheI (66) using AgeI and NheI restriction enzyme sites. For the *flhDC* expression strains, the nucleotide sequences *flhDC* (amplified using oKL277 and oKL278) was similarly inserted into pBBR1-NheI. A gBlock containing the *lac* promoter (Integrated DNA Technologies, Coralville, IA) was introduced upstream of the coding region using SLiCE (69).

### LPS extraction and analysis

Cells were grown on overnight at 37°C on swarm-permissive CM55 (Oxoid, Hampshire, UK) agar with antibiotics as needed and harvested with LB. LPS from cells was extracted using an LPS Extraction Kit according to manufacturer’s instructions (iNtRON Biotechnology Inc, Sangdaewon Seongnam, Gyeonggi, Korea). Extracts were resuspended in 10mM Tris, pH 8.0 buffer and run on a 12% SDS-PAGE gel. Gels were stained with a modified silver stain protocol (45).

### Halo assays (MIC determination)

Cultures were top-spread on LSW-medium and allowed to sit on benchtop until surface appeared dry (a couple of hours). 6 mm sterile filter disks were placed onto plates and soaked with 10 µL of dilutions containing Polymyxin B (Sigma Aldrich, St Louis, MO), gentamycin (Calbiochem, San Diego, CA), or kanamycin (Corning, Corning, NY) in water. A water-alone control was included. Once filter disks dried (a couple of hours), plates were incubated at 37°C overnight and imaged.

### Bile salts and SDS resistance assays

Cultures were grown overnight, normalized to OD 1.0, and serial diluted 10-fold. 1 µL spots of 10^-1^ to 10^-8^ dilutions of each strain (in technical triplicate) were inoculated onto LSW-plates containing bile salts (Sigma Aldrich, St Louis, MO) or sodium dodecyl sulfate (SDS) (Sigma Aldrich, St Louis, MO) (filter sterile, added to medium post-autoclaving). Plates were incubated overnight at 37°C.

### Carbenicillin sensitivity assay

Cells were harvested from swarm permissive medium using LB broth and spread onto 1 mm CM55 (Oxoid, Hampshire, UK) agar pads containing 10 µg/mL carbenicillin (Corning, Corning, NY). Cells were imaged after 1, 2, or 4 hours on surface. Three biological replicates were analyzed.

### Microscopy

Microscopy was performed as previously described (68). Briefly, CM55 (Oxoid, Hampshire, UK) agar pads, supplemented as needed with 35 µg/mL kanamycin for plasmid retention, were inoculated from overnight stationary cultures and incubated at 37°C in a modified humidity chamber. Pads were imaged using a Leica DM5500B (Leica Microsystems, Buffalo Grove, IL) and a CoolSnap HQ2 cooled CCD camera (Photometrics, Tuscon, AZ). MetaMorph version 7.8.0.0 (Molecular Devices, Sunnyvale, CA) was used for image acquisition. Images were analyzed using FIJI (72) (National Institutes of Health, USA); where indicated, images were subjected to background subtraction equally across entire image. Where indicated, cells were stained with 25 µM TMA-DPH (Invitrogen, Carlsbad, CA), (max excitation 355 nm; max emission 430 nm) imaged in the DAPI channel using an A4 filter cube (excitation 360/40 nm; emission 470/40 nm) (Leica Microsystems, Buffalo Grove, IL). Venus (max excitation 515 nm; max emission 528 nm) was visualized in the GFP channel using a GFP ET filter cube (excitation 470/40 nm; emission 525/50 nm) (Leica Microsystems, Buffalo Grove, IL). Fluorescence intensity and exposure times for each fluorescence channel were equivalent across all fluorescence microscopy experiments. Fluorescence due to Venus was not quantified and was visible due to the overlapping excitation and emission spectra with the GFP ET filter cube.

### Transcriptional analysis

Strains were grown on CM55 (Oxoid, Hampshire, UK) plates at 37°C. For wild-type samples, colonies were inoculated on CM55 (Oxoid, Hampshire) agar and incubated overnight for swarm development. The presence of short, non-motile cells in consolidation phase was confirmed by light microscopy. Wild-type cells from the swarm edge were then harvested by scraping with a plastic loop into 1 ml of RNA Protect solution (Qiagen, Venlo, Netherlands). Samples of the Δ*rffG* strain were harvested after overnight incubation by scraping whole colonies into 1 ml RNA Protect solution. Total RNA was isolated using an RNeasy Mini kit (Qiagen, Venlo, Netherlands) according to the manufacturer’s instructions. RNA purity was measured using an Agilent 2200 Tapestation (Agilent, Santa Clara, CA). To enrich mRNA, rRNA was digested using terminator 5’ phosphate dependent exonuclease (Illumina, San Diego, CA) according to the manufacturer’s instructions and purified by phenol-chloroform extraction (73). cDNA libraries were prepared from mRNA-enriched RNA samples using an NEBNext Ultra RNA library prep kit (New England Biolabs, Ipswich, MA) according to the manufacturer’s instructions. Libraries were sequenced on an Illumina NextSeq 2500 instrument with 250-basepair single-end reads at the Harvard University Bauer Core. Sequences were matched to the BB2000 reference genome (accession number CP004022) using Tophat 2 (74). Differential expression data were generated using the Cufflinks RNA-Seq analysis suite (75) run on the Harvard Odyssey cluster, courtesy of the Harvard University Research Computing Group. Data were analyzed using the CummeRbund package for R and Microsoft Excel (75). Bioinformatics information was derived from KEGG (76). The data in this paper represent the combined analysis of two independent biological repeats.

## Acknowledgements

We would like to thank Drs. Enrique Balleza and Philippe Cluzel for the kind gift of the Venus construct and members of the Gibbs, D’Souza, Gaudet, and Losick labs for thoughtful discussion and feedback. Drs. Doug Richardson, Sven Terclavers, and Sebastian Gliem assisted with imaging on the Cell Observer. We would like to thank the staff of the Bauer Core at Harvard University for assistance with RNA-Seq and Maria Ericsson of the Harvard Medical School Electron Microscopy facility for assistance with TEM (Supplemental Information). This research was funded by a Smith Family Fellowship in Science and Engineering, a Simmons Family Award for the Harvard Center for Biological Imaging, the David and Lucile Packard Foundation, the Merck Family Fund, and Harvard University.

## Author Contributions

KL and KAG conceived and coordinated the study and wrote the paper. KL performed all experiments, except the RNA-Seq which was performed by MJT. All authors edited the paper.

